# Precise timing of sensory modulations coupled to eye movements during active vision

**DOI:** 10.1101/144477

**Authors:** Christ Devia, Rodrigo Montefusco-Siegmund, José Ignacio Egaña, Pedro E. Maldonado

**Affiliations:** Programa de Fisiología y Biofísica, Facultad de Medicina, Universidad de Chile.; Biomedical Neuroscience Institute, Universidad de Chile.; Departamento de Anestesiología y Reanimación, Facultad de Medicina, Universidad de Chile.; Escuela de Psicología, Pontificia Universidad Católica de Chile, Santiago, Chile.

## Abstract

Perception is the result of ongoing brain activity combined with sensory stimuli. In natural vision, changes in the visual input typically occur as the result of self-initiated eye movements. Nonetheless, in most studies, stimuli are flashed, and natural eye movements are avoided or restricted. As a consequence, the neural sensory processing associated with active vision is poorly understood. Here, we show that occipital event-related potentials (ERP) to eye movements during free exploration of natural images exhibited different amplitudes, time course and motor dependency than that from the same flashed stimuli. We found that the ERP to visual fixations doubles in P1 magnitude and does not show a late component, which is classically seen with flashed stimuli^1,2^. In addition, we discovered that the ERP to the saccade onset was as large as the ERP to fixations onset, with an early component that preceded the visual input, suggesting that a motor modulation was associated with the saccades^3^. Furthermore, the use of different visual scenes revealed that both the ERP amplitude and time course were dependent on the type of image explored. Our results demonstrated that during active vision, the nervous system engages a mechanism of sensory modulation that is precisely timed to the self-initiated stimulus changes. This mechanism could help coordinate neural activity across different cortical areas and, by extension, serve as a general mechanism for the global coordination of neural networks.

## Results

Event-related potentials (ERP) obtained from electroencephalographic (EEG) signals have been one of the fundamental tools for studying the sensory and cognitive processes in the brain. Characteristically, these signals represent the average activity resulting from the repetitive presentation of a stimulus. In most vision studies to date, the stimuli are flashed in front of subjects, a situation that contrasts with natural vision where subjects, by moving their eyes, self-initiate changes in the visual input. Although saccade (saccRP), fixation (fixRP), and image (imgRP) related potentials have been independently explored^1,4-6^, the neuronal processes that underlie these different responses are not well understood. We hypothesize that a direct comparison of these responses would reveal distinct mechanisms involved in passive and active vision.

In this work, we assessed the magnitude of the modulation by self-driven eye movements on the activity in the early visual areas in human subjects. During each trial, the subjects freely explored pictures (Fig. 1a) while their eye movements and EEG activity were recorded (Fig. 1b). The subjects (n = 16) freely explored natural scenes (NS) and 6 other control categories (pink noise, white noise, and white, grey, and black images; Fig. 1a).

**Figure 1.**
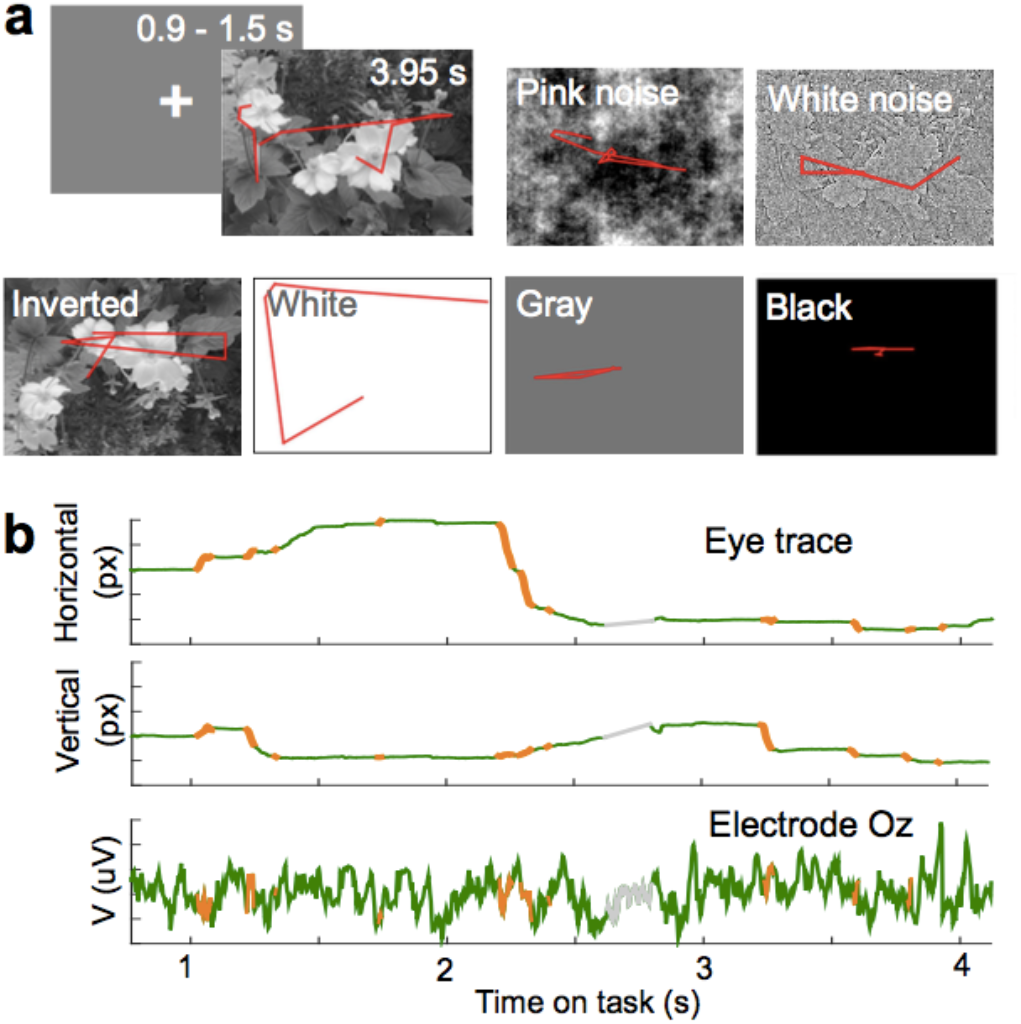
Free-viewing task, eye movements and EEG recordings. **a**. The subjects freely explored images from 7 categories (natural scenes, pink noise, white noise, inverted scenes, and white, grey, and black images). Over an example image is the eye trace (red) of one subject in all the image categories tested. This example image was not part of the database. **b**. The time course of the horizontal (Hor) and vertical (Ver) component of the eye movement, and simultaneous EEG recording (bottom) for the eye trace in the NS of **a**. Epochs drawn in green, orange, and grey show fixation, saccade, and blinks, respectively.

We first compared the event related potential to the fixation onset with the potential to image onset (fixRP *vs*. imgRP). The fixRP exhibited a significantly larger amplitude and a shorter latency than the imgRP (Fig. 2a), with its maximum amplitude observed in the occipital electrodes (O1, Oz, and O2; see the topography plot in Fig. 2a). Therefore, all following analyses were performed on the average of those three electrodes. A prominent positive first peak characterized the fixRP, which was preceded and followed by brief and small negative potentials. The fixRP waveform was consistent with high frequency power from the electrocorticogram recordings in the early visual areas in humans^1^ (mainly V1, V2, and V3) and the firing rate in non-human primates^2^ (V1, V2, and V4) during free viewing tasks, suggesting that the EEG probes over the occipital areas were mainly recording the firing rate on the visual cortex. The positive component (fixP1, latency 88.10 ± 8 ms and mean peak amplitude 5.48 ± 2.91 μV) was visible on each subject, and its waveform was consistent between the subjects, showing only amplitude modulation. An additional feature of the fixRP was the negative potential previous to the fixP1 that started even before the fixation (onset around -76.04 ± 33.65 ms, min amplitude -1.34 ± 0.77 μV at 12.71 ± 18.94 ms; Fig. 2a). It has been shown that this negative potential is linearly modulated by the contrast between successive fixations and nonlinearly by the duration of the fixation^1^. Because this potential begins before fixation, it suggests that the brain events related to current stimulus processing on the sensory surface started during the saccade. The second negative potential that followed fixP1 had even less amplitude (-1.46 μV at 140 ms), and its amplitude is modulated by the fixation duration^1^. As previously reported^6^, the image related potential (imgRP) exhibited two positive peaks, where the first peak had a smaller amplitude than the fixRP (the imgP1 mean peak amplitude on NS was 2.47 ± 2.87 μV and the peak latency was 105.40 ± 15.6 ms), and the second potential had a longer duration and amplitude similar to the fixP1 (the imgP2 mean area under the curve was 299.01 ± 181.12 μVms and the peak latency was approximately 210 ms). The imgP1 amplitude correlates with the stimulus contrast^3^, and its origin is probably in the dorsal extrastriate cortex^7^. The second potential on the imgRP, the imgP2, has been proposed to correspond to the neural correlate of the gist of the scene^6,8^, where the recognition of the main features of the image may occur^9^. The time course of the fixRP and the imgRP substantially differed. The fixP1 peaked significantly earlier than the imgP1 (Fig. 2a, left insert), and despite the large luminance changes that occur at the image onset, the fixP1 had a significantly larger amplitude than the imgP1 (Fig. 2a, right insert). We demonstrate that self-triggered changes in the visual stimulus, generated by the exploratory eye-movements, correlate with a modulation of EEG signals in the human visual cortex that is stronger than the activity evoked by the onset of the NS image.

**Figure 2.**
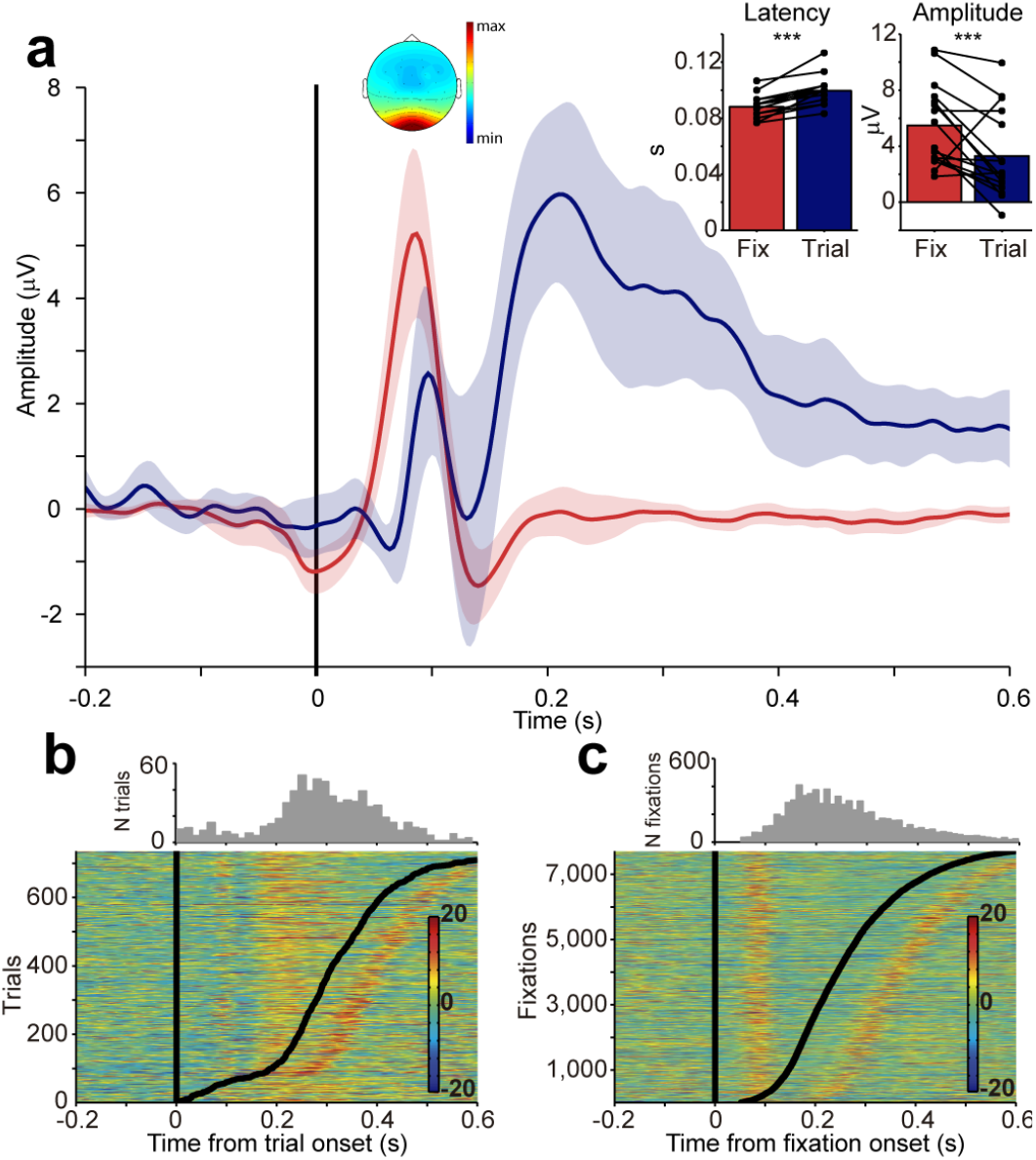
Fixation-related potentials exhibit different amplitudes and time courses than image-evoked occipital potentials on natural scenes. **a**. The occipital evoked response for NS aligned to the fixation onset (red) and image onset (blue). The shadowing depicts confidence intervals. The occipital distribution of fixP1 is over the scalp topography (average over 43 to 117 ms). Left panel, the mean latency of fixP1 and imgP1 (latency of fixP1 *vs*. latency of imgP1, t_15_ = -7.84, P = 1.1x10^−6^, paired *t*-test). Right panel, mean amplitude of fixP1 and imgP1 across subjects (peak amplitude of fixP1 versus imgP1, t_15_ = 3.12, P = 0.007, paired *t*-test). The dots represent each subject during each condition connected by lines. **b**. Top: The histogram shows the distribution of latencies to the first fixation on the image. Bins of 15 ms. Bottom: The single-trials from imgRP aligned to the image onset and sorted by the first fixation latency (black curved line). **c**. Top: The histogram shows the distribution of the next fixation latency. Bins of 11 ms. Bottom: The single-trials activity of fixRP aligned to the fixation onset and sorted by the next fixation latency (black curved line).

Previous work on non-human primates showed that the first eye movement is typically suppressed until 200 ms after image onset^10^. Consequently, we investigated the time of occurrence of the first fixations after image onset and examined its contribution to imgP2. Specifically, we determined whether this signal corresponded solely to evoked activity by the image or was combined with a putative response to the first eye movement after image onset. As expected, we found that from the 735 first fixations examined, 15.24% occurred in the first 200 ms after image onset, while 64.63% first fixations occurred during the period 200-400 ms after image onset (histograms on Fig. 2b, c). All trials were then sorted by the latency of the first fixation (Fig. 2b). We found that the imgP2 component precedes the increment in the number of first fixations (>200 ms). Moreover, the imgP2 positive potential was clearly distinguishable from the subsequent fixP1, and it exhibited a longer duration (the black curved line depicts the fixation onset in Fig. 2b). In contrast, when the single-trial of fixRP was sorted by the latency to the next fixation (Fig. 2c), the fixP1 potential (the black curved line in Fig. 2c) exhibited a shorter and precise time course, and we did not observe a second peak regardless of the fixation duration. These results suggest that the fixRP and the imgRP originate on different brain processes, where the fixRP is locked to the self-paced eye movement, while imgRP is mostly evoked by stimulus onset associated with gist and other high-level cognitive processes ^6,11,12^.

A key feature of active vision is that the self-initiation of a saccade results in an ensuing and timely visual stimulation. In the primary visual cortex, it has been shown that the local field potential displays greater amplitude when the signals are aligned to the onset of the saccades, suggesting an influence of these motor actions in V1 activation^4^. Here, we found that the saccade related potential (saccRP) presented a negative potential that started before saccade onset, followed by a positive potential that was larger in amplitude than the fixP1 and that it had a nonlinear modulation by the saccade amplitude. The saccRP (Fig. 3a) had a negative component that started before saccade onset, which we termed Nz (Negative-zero). This component was followed by a positive potential (saccP1) that peaked after saccade onset and ended in another negative deflection. The shape of the saccRP, especially the saccP1 and the following negativity, resembles the event related potential from the electrocorticogram recordings in areas V1 and V4^13^ and the firing rate on V1^5^ in monkeys. Our data revealed that the Nz was also visible in the fixRP but had significantly reduced amplitude than in the saccRP (-1.67 ± 0.86 μV at 48.75 ± 15.09 ms; Fig. 3a and inset). The duration of the Nz, from its onset until the minimum value before P1, was significantly longer than in the fixRP (Fig. 3a, right inset). The Nz also showed a spike potential coincident with the saccade initiation and duration, and thus, likely originated in the extraocular muscles that move the eyeball^14–16^. This shows that the spike potential captured the muscle twitch artefact and that this muscle activity was distinguishable from the occipital neuronal processes. The amplitude of the saccP1 (6.02 ± 3.23 μV at 120±8.5 ms) was significantly larger than that of the fixP1 (Fig. 3a), and peaked 36 ms after the fixP1, a time lapse coincident with the median duration of saccades^17^ (here, the median saccade duration was 31.98 ms, with 10 ms and 60 ms at 5% and 95% of the distribution, respectively). These results for the Nz and saccP1 show that brain activity from occipital areas is larger in amplitude and better timed to the saccade onset rather than to the fixation onset. We conclude that saccades are instrumental in modulating the visual processing of the subsequent foveated visual patches. For short saccade lengths, the amplitude of the saccP1 increased with the amplitude of the movement (Fig. 3b), showing a linear correlation for saccades smaller than 2.5° (Fig. 3c), which is intriguingly within the fovea size range (the foveal diameter is approximately 5°). However, for saccades larger than the fovea radius, the saccP1 amplitude seems to saturate regardless of the saccade length (Fig. 3c). These findings demonstrate that this component of the response cannot be of purely motor origin because otherwise, its amplitude should keep growing linearly with the saccade length. Additionally, the relationships between saccade length, foveal size, and saccP1 support previous studies that reported that humans and monkeys employ ambient and focal modes of visual exploration while freely viewing natural scenes^18,19^. Finally, the negative potential that came after the saccP1 was similar between the saccRP and fixRP. This potential is coincident with the suppression of the firing rate in V1 for small saccades^5^, which is absent when the movement is externally induced instead of self-initiated. Together, these results demonstrate that during free viewing of NS, the processing of the upcoming stimulus in the early visual areas is timed to the saccade onset and that saccade-related brain activity contributes to the processing of visual images.

**Figure 3.**
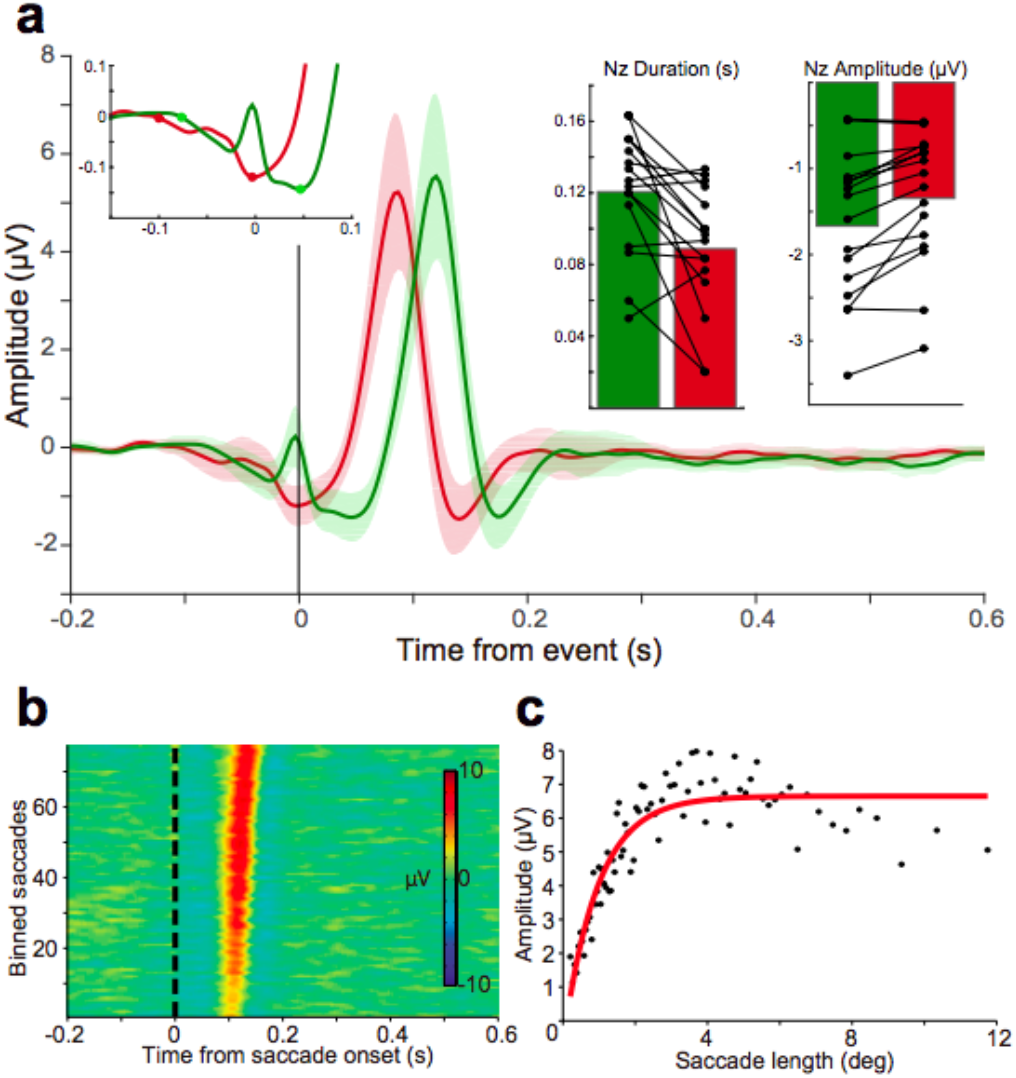
The saccade related potential (saccRP) is larger than the fixRP and exhibits an early component that is differentially modulated by short and long saccades. **a** The saccRP (green) and fixRP (red) are plotted on the same time axes. The saccP1 was bigger than the fixP1 (t_15_ = -2.75, P = 0.02, paired *t*-test). Time zero is the event onset, saccade or fixation, respectively. The shaded area is the confidence interval. Inset: The details on the Nz, the negative potential that started before the saccade onset. The dots over the curves denote the landmarks used to measure Nz duration. Inset, left: The bar plot shows the duration of the Nz on the saccRP and fixRP (saccRP 120.63 ± 33.82 ms, Nz_fix_ *vs*. Nz_sacc_ t_15_ = -3.47, P = 0.003, paired *t*-test). Inset, right: The Nz has a larger amplitude on the saccRP than on the fixRP (Nz amplitude on the fixRP *versus* the saccRP, t_15_ = -4.50, P = 4.26x10^−4^, paired *t*-test). The dots represent each subject on each condition connected by lines. **b**. The mean EEG activity on bins based on saccade length. Each bin is the average of 100 trials previously sorted by saccade length. **c**. The mean saccP1 and mean saccade length on binned data from b (Correlation for saccade length ≤ 2.5° was ρ = 0.93, P = 0; for longer saccades ρ = -0.22, P = 0.19 Spearman correlation).

Previous studies have shown that the fixRP is modulated by the luminance contrast between two fixation points^20^. To further investigate how low-level features of visual scenes modulate the saccRP, we measured the saccRP in plain (white, black and grey images) and textured images (pink noise, white noise, and inverted images, see the Methods section and Fig. 1a) and found that the saccRP also depended on the content of the image (Fig. 4a). The saccRP on the plain images showed a smaller and delayed development of a positive deflection approximately 150 ms after saccade onset, which is in contrast with the well-defined saccP1 on NS. As a group, the textured images showed a significantly larger amplitude of Nz compared with the plain images. Similarly, the saccP1 amplitude was significantly increased in the textured images compared with the plain images. Specifically, the saccRP in the plain images had a reduced Nz (-0.57 ± 0.44; white -0.45 ± 0.40 μV, black -0.46 ± 0.43 μV, grey -0.79 ± 0.52 μV), and the saccP1 was almost absent (white 0.60 ± 0.99 μV, black 0.50 ± 0.91 μV, grey 0.69 ± 0.83 μV). These reduced potentials on plain images are in accordance with a general reduction in the firing rate on V1 when a grey screen is on the receptive field of monkeys^5^. In contrast, for textured images (pink noise, white noise, and inverted images), we found that the Nz and the saccP1 were conserved and had similar magnitudes to NS potentials. The Nz potential was similar across the textured categories (NS -1.62 ± 0.73 μV, pink noise -1.39 ± 0.71 μV, white noise -1.45 ± 0.81 μV, inverted images -1.32 ± 0.67 μV). These large differences in Nz signals cannot be attributed to motor modulation because the subjects still produce eye movements in plain as in textured images. The saccP1 amplitude was similar between the NS (6.02 ± 3.23 μV), pink noise (5.68 ± 3.29 μV), and inverted images (5.57 ± 3.02 μV; Fig. 4a inset). Notably, the white noise images elicited the largest saccP1 amplitude^21^ (7.11 ± 3.84 μV), even larger than the NS images. This latter result can be explained, in part, by the change in luminance between two successive fixations^20^. However, it can also be interpreted as a decrement in the predictability of the content of the next fixation^22^. Further examination of the saccP1 and its relation with saccade length supports this latter hypothesis, where the linear relation between the saccP1 and saccade length exhibited a steeper slope for the white noise images than the NS images. This would also explain our findings of significantly smaller saccP1 amplitudes for the plain images, given that there is no luminance contrast between successive fixations and full predictability of the content of the next fixation. A putative explanation to conciliate the reduction in both Nz and sacP1 in the plain images would suggest that as quickly as the gist of the image is acquired after image onset, a distinct visual process ensues, which would depend on the nature of the gist. As a result, on plain image presentations, the motor-related modulation observed in the textured images may not be engaged, as subjects quickly perceive a uniform visual scene. However, for the textured images, each eye movement resulted in a new stimulus to early visual areas that needed to be processed. Thus, motor-related modulation becomes an important part of the visual processing of each fixation. This conjecture is consistent with our finding that the Nz and saccP1 on the plain images were not due to some lack of general response to the stimulus. Indeed, when we examined the response to the onset of the images, we verified that plain images were able to produce the expected imgP1 component and a weak imgP2, with similar timing to imgRP on NS (Fig. 4b). There were significant differences for the imgP1 between the plain and NS categories (NS, white, grey, and black). The transition from the fixation cross (grey background during the inter-trial interval; Fig. 1a) to the grey images evoked an imgP1 of smaller amplitude than in the NS (grey 0.74 ± 1.45 μV and NS 2.47 ± 2.87 μV), and for white and black images, the imgP1 was larger than in the NS (3.93 ± 2.69 μV and 4.42 ± 3.69 μV, respectively), but not significantly. The imgP1 was significantly larger for the white and black images than the grey. These results, as with the NS images (Fig. 2a), further indicate that the imgP1 was mainly modulated by the difference in luminance between the current and the previous image^3^. The late component, imgP2, also had significant differences between categories (Fig. 4b). The imgP2 was similar between the plain images (white 82.53 ± 104.84 μV, black 95.91 ± 90.54 μV, and grey 83.52 ± 74.89 μV), and significantly smaller than the NS (299.01 ± 181.12 μV). These results further support the interpretation of imgP2 as the correlate of the gist of the scene^6^.

**Figure 4.**
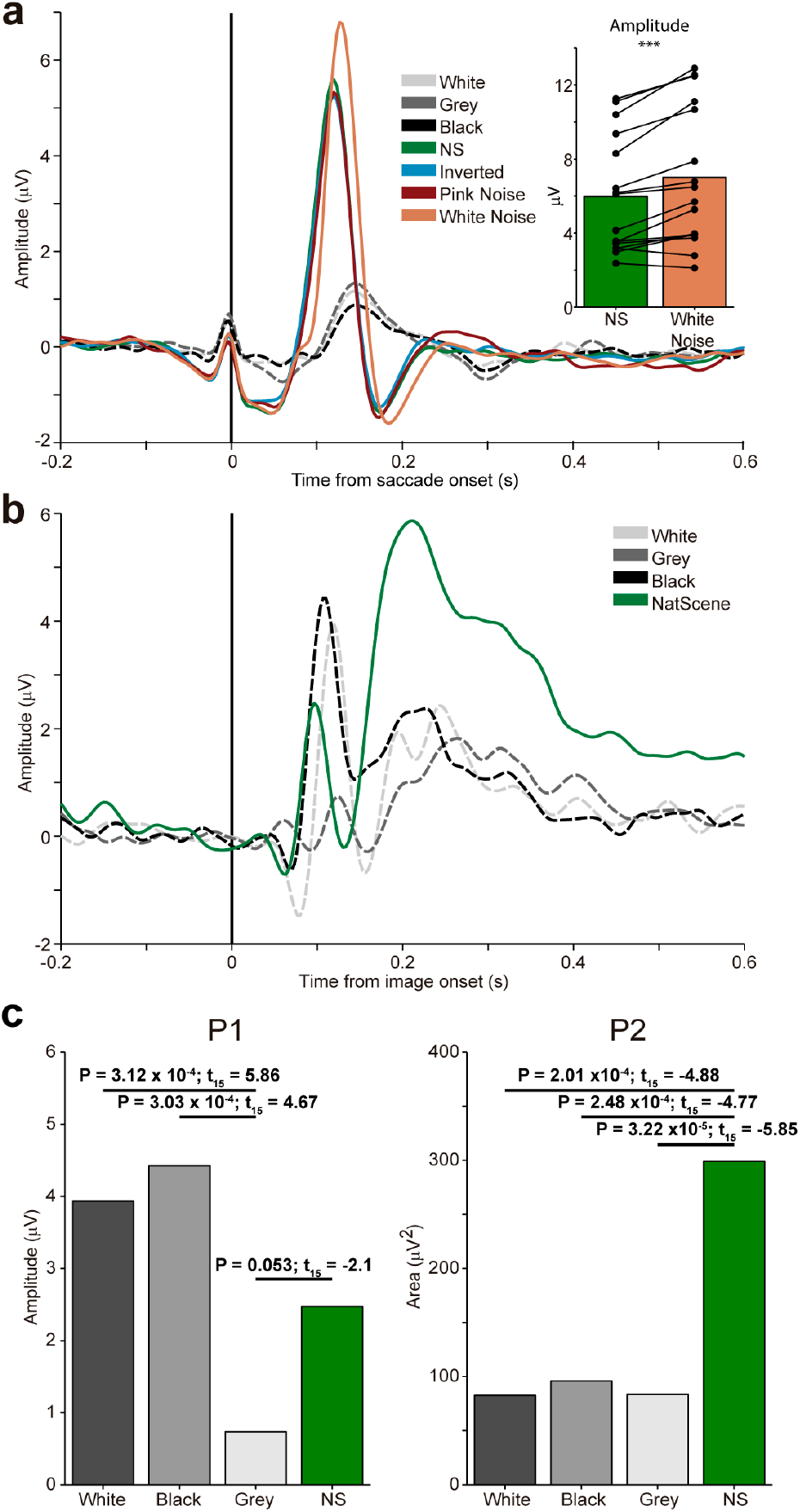
The saccRP and imgRP on plain and complex images. **a**. The saccRP on plain (white, grey, and black) and complex images (NS, inverted, pink noise, and white noise; Nz from all content images versus all plain images, t_15_ = -4.84, P = 2.18x10^−4^, paired *t*-test; the same for saccP1, t_15_ = 7.55, P = 1.74x10^−6^, paired ŕ-test). Inset, the white noise saccP1 compared with the NS (t_15_ = -5.01, P = 1.53x10^−4^, paired *t*-test). **b**. The imgRP on the NS and plain images (imgP1, F_3,45_ = 7.75, P = 2.79x10^−4^; imgP2, F_3,45_ = 21.60, P = 8.17x10^−9^, one-way ANOVA repeated measures). **c**. Left: A bar plot of imgP1 amplitude in the NS and plain images. No significant differences between NS and grey (P = 0.46) or between white or black images and NS (P = 0.16 and P = 0.09, respectively). Right: A bar plot of the imgP2 amplitude in the NS and plain images. (P = 0.53 white vs. black, P = 0.96 white vs. grey, P = 0.50 black vs. grey). A paired *t*-test was carried out for all bar plots.

## CONCLUSIONS

Our results show that during self-paced image exploration, the activity in the brain occipital areas is time-modulated to the saccade onset rather than to the fixation onset. The evoked activity to the saccade onset (saccRP) has increased amplitude compared to the activity evoked by the fixation onset. Moreover, these potentials were greater in magnitude than the response to image onset. Our results provide additional evidence of the role of internally generated motor signals in the modulation of neural activity in early visual areas^4^ and suggest that self-initiated sensory activity triggers different neural processes than those triggered from externally imposed stimuli. Another important finding was that the amplitude of the saccP1 for the natural scenes is linearly correlated with saccade length that fall within the fovea radius. This result fits well with the idea of a local-global strategy working during free visual exploration^19,23^. Additionally, we found that the saccRP is almost absent for images without texture (plain images), even though subjects still move their eyes. A possible explanation for this latter result is that the internally generated motor signal is not engaged during exploration of the plain images. This implies that, shortly after image presentation, the perceptual process categorizes images as plain or textured. As expected, the imgP2 depends on the class of visual stimulus and was diminished for non-textured images, supporting imgP2’s role in the assessment of scene content. This finding extends previous reports proposing that the imgP2 is the neural correlate of the gist of an image^6^.

Altogether, our results suggest that during image exploration, two different modes that are mutually exclusive can be engaged. A sensory motor modulation over early visual cortex acts when the explored scene has textures, but it is disengaged when an image is categorized as plain. The findings from this research have important implications for classical paradigms in visual neuroscience, where stimuli are flashed to the subjects in an externally paced frequency and highlights the need to complement the current body of literature with free exploration paradigms, where intrinsic brain processes are exposed. We should ask not what the brain can do but what the brain actually does.

## Methods

### Subjects

A total of 16 subjects participated in this experiment (5 female); 13 were right-handed, 8 had right ocular dominance, and their average age was 31.41 ± 7.17 years. All have normal or corrected-to-normal vision. All subjects were volunteers and accepted to participate through a written informed consent; the consent form and all experimental protocols were approved by the Ethics Committee for Research in Humans of the Faculty of Medicine from Universidad de Chile. All recordings were performed within a 32-day period.

### Stimuli and task

A total of 46 natural images, from the International Affective Picture System (IAPS)^24^, were selected based on their valence and arousal value, with all ranging in the middle values of the database (mean valence 6.62 ± 1.09; mean arousal 4.08 ± 0.96). The selected images were gamma corrected for the screen used to present the stimuli, and these images constituted the NS set. For each of the NS images, we constructed 4 control images: pink noise, white noise, inverted, and grey. The pink noise and white noise images were created considering 2 parameters of the NS – its power spectrum and its phase. The pink noise images had the same spectral power as that of the NS, with the phase from a scrambled version of the original NS. As a complement, the white noise images were created having a flattened power spectrum of the scrambled version of the NS and the phase content of the original NS. The inverted images were created as a high-level control, where they were the original NS upside down with the bottom part at the top the image. The grey images had the same average luminance as that of the original NS. Finally, the white and black images were also created as controls of the luminance, and they had the maximum ([255,255,255] RGB) and minimum luminance ([0,0,0] RGB), respectively. The subjects’ eyes were 70 cm from the screen. The screen size was 1920 (width) x 1080 (height) pixels, and the mean 32 pixels per cm was equivalent to 39.38 pixels per visual degree. The image size was 1024 x 768 pixels, and it was centred on the screen. The remaining borders in the screen were a plain grey colour ([173,173,173] RGB). The subjects were seated in a dark room with their chin rested. We first recorded a couple of minutes of the EEG while the subjects rested (data not analysed here). Then, they performed the main task where they were instructed to explore the images freely, as there would be questions about their content at the end. Here, a total of 322 images divided into 2 blocks were presented. Finally, on a third task, they observed 54 images, where 8 were not part of the main task, and they were asked whether they remembered seeing the image.

### Recordings

We recorded the brain activity using 32 electrodes (ActiveTwo, BioSemi, Amsterdam, The Netherlands) displayed as in the International 10-20 system. In addition, eight electrodes, one on each mastoid for re-referencing, while the other six recorded the electro-oculogram (EOG), were placed above, below, and on the outer canthi of each eye. The EEG and EOG were recorded at 2048 Hz. The gaze position and pupil diameter were recorded with a chin rest eye tracking system at 1000 Hz (EyeLink 1000, SR-Research, ON, Canada).

### Eye movement analysis

For all analyses of eye movements, we considered only the data obtained for the right eye. The saccades and fixations were automatically detected based on the velocity (30 °/s) and acceleration threshold (8000 °/s^2^). The saccades longer than 5 ms and smaller than 100 ms and the fixations between 50 and 600 ms were considered for further processing. The blinks were defined as the absence of pupil data. The saccades that were actually blinks were discarded (a median of 9.32% of the events were blinks).

### EEG and ERP analyses

The EEG processing was done in Matlab (release 2012b; The MathWorks, Inc., Natick, Massachusetts, United States) using the toolbox FieldTrip (release 20140412)^25^. The recordings were re-referenced to the average of the mastoids. Then, we performed an ICA decomposition of the data^26^ and visually compared the EOG channels with the ICA components to discard the ICA components that had ocular content from the remaining components, and we reconstructed the signal. Then, the data were decimated to a sample rate of 300 Hz. We applied a band-pass filter (Butterworth) between 0.5 and 30 Hz. The epochs were defined 1 second before the event onset until 1 second after. We used the period [-200, -100] ms as baseline for fixRP and saccRP and between [-200, 0] ms for imagRP. We studied the average event-related potentials from the occipital electrodes (O1, Oz, and O2) time-locked to fixation, saccade, or image onset, respectively. For each subject, we assessed the peak amplitude of these potentials, while for imgP2, we considered the area under the curve. We used the area under the curve because it was not possible to detect a clear maximum for the imgP2 of each subject^27^. The onset of Nz was defined as the first point between [-100,+100] ms, where three consecutive samples were below the baseline. The offset was the point where the saccRP reached a minimum, just before the positive deflection of saccP1 began.

### Statistics

We measured the significant differences for fixRP *versus* imgRP, and fixRP *versus* saccRP with paired *t*-tests. The subjects were the independent samples, and we reported the *t*-statistics, with its degrees of freedom, and P-value. We applied the test over the maximum or minimum amplitude of the potentials (for sacc/fix/imgP1 and sacc/fix Nz, respectively) and the area under the curve for imgP2. To assess the significant differences across image categories, we performed an ANOVA of repeated measures and reported the F-statistic of times (image categories), with the degrees of freedom of the times and error (F_dfTimes, dfError_), along with its P-value.

## Acknowledgements

This work was made possible in part by a grant from CONICYT, FONDECYT/Postdoctorado 3140306 to CD and by ICM-P09-015F.

## References

1. Podvalny, E., Yeagle, E., Mégevand, P., Sarid, N. & Harel, M. Invariant Temporal Dynamics Underlie Perceptual Stability in Human Visual Cortex. Current Biology (2016).

2. Gallant, J. L., Connor, C. E. & Van Essen, D. C. Neural activity in areas V1, V2 and V4 during free viewing of natural scenes compared to controlled viewing. Neuroreport (1998).

3. Johannes, S., Münte, T. F., Heinze, H. J. & Mangun, G. R. Luminance and spatial attention effects on early visual processing. Brain Res Cogn Brain Res 2, 189–205 (1995).

4. Ito, J., Maldonado, P. & Grün, S. Cross-frequency interaction of the eye-movement related LFP signals in V1 of freely viewing monkeys. Front Syst Neurosci 7, 1 (2013).

5. Troncoso, X. G. et al. V1 neurons respond differently to object motion versus motion from eye movements. Nat Commun 6, 8114 (2015).

6. Harel, A., Groen, I. I. A., Kravitz, D. J., Deouell, L. Y. & Baker, C. I. The Temporal Dynamics of Scene Processing: A Multifaceted EEG Investigation. eNeuro 3, (2016).

7. Di Russo, F., Martínez, A., Sereno, M. I., Pitzalis, S. & Hillyard, S. A. Cortical sources of the early components of the visual evoked potential. Hum Brain Mapp 15, 95–111 (2002).

8. Thorpe, S., Fize, D. & Marlot, C. Speed of processing in the human visual system. Nature (1996).

9. Oliva, A. & Torralba, A. Building the gist of a scene: the role of global image features in recognition. Prog. Brain Res. 155, 23–36 (2006).

10. Bartlett, A. M., Ovaysikia, S., Logothetis, N. K. & Hoffman, K. L. Saccades during object viewing modulate oscillatory phase in the superior temporal sulcus. J. Neurosci. 31, 18423–18432 (2011).

11. Devillez, H., Guyader, N. & Guérin-Dugué, A. An eye fixation-related potentials analysis of the P300 potential for fixations onto a target object when exploring natural scenes. Journal of Vision 15, 20 (2015).

12. Villena-González, M., López, V. & Rodriguez, E. Orienting attention to visual or verbal/auditory imagery differentially impairs the processing of visual stimuli. Neuroimage 132, 71–78 (2016).

13. Brunet, N. et al. Visual cortical gamma-band activity during free viewing of natural images. Cereb. Cortex 25, 918–926 (2015).

14. Kaunitz, L. N. et al. Looking for a face in the crowd: fixation-related potentials in an eye-movement visual search task. Neuroimage 89, 297–305 (2014).

15. Ioannides, A. A., Fenwick, P. B. C. & Liu, L. Widely distributed magnetoencephalography spikes related to the planning and execution of human saccades. J. Neurosci. 25, 7950–7967 (2005).

16. Carl, C., Açık, A., König, P., Engel, A. K. & Hipp, J. F. The saccadic spike artifact in MEG. Neuroimage 59, 1657–1667 (2012).

17. Bahill, A. T., Clark, M. R. & Stark, L. The main sequence, a tool for studying human eye movements. Mathematical Biosciences (1975).

18. Borji, A. & Itti, L. State-of-the-art in visual attention modeling. IEEE Trans Pattern Anal Mach Intell 35, 185–207 (2013).

19. Ito, J. et al. Switch from ambient to focal processing mode explains the dynamics of free viewing eye movements. Sci Rep 7, 1082 (2017).

20. Ossandón, J. P., Helo, A. V., Montefusco-Siegmund, R. & Maldonado, P. E. Superposition model predicts EEG occipital activity during free viewing of natural scenes. J. Neurosci. 30, 4787–4795 (2010).

21. Katz, L. N., Yates, J. L., Pillow, J. W. & Huk, A. C. Dissociated functional significance of decision-related activity in the primate dorsal stream. Nature 535, 285–288 (2016).

22. Rao, R. P. & Ballard, D. H. Predictive coding in the visual cortex: a functional interpretation of some extra-classical receptive-field effects. Nat. Neurosci. 2, 79–87 (1999).

23. Berger, D. et al. Viewing strategy of Cebus monkeys during free exploration of natural images. Brain Research 1434, 34–46 (2012).

24. Lang, P. J., Bradley, M. M. & Cuthbert, B. N. International affective picture system (IAPS): Affective ratings of pictures and instruction manual. (Technical report A-8, 2008).

25. Oostenveld, R., Fries, P., Maris, E. & Schoffelen, J.-M. FieldTrip: Open source software for advanced analysis of MEG, EEG, and invasive electrophysiological data. Comput Intell Neurosci 2011, 156869 (2011).

26. Comon, P. Independent component analysis, a new concept? Signal processing (1994).

27. Picton, T. W. et al. Guidelines for using human event-related potentials to study cognition: recording standards and publication criteria. Psychophysiology 37, 127–152 (2000).

